## Supplementary Material for "Overlapping neural representations for the position of visible and imagined objects"

**S1. Analysis of correct trials only**

Performance on the task was high (*M* > 80%). We chose to analyse all trials in the decoding analyses. However, it is possible that on incorrect sequences participants did not track the stimulus correctly and would have neural responses consistent with the wrong position, affecting the decoding. To assess position-related information on correct trials, the time-resolved decoding analysis was performed again by excluding incorrect sequences from the test set. Number of trials included are listed in Table S1. As can be seen in Figure S1, the temporal dynamics of visible and imagined stimulus position are very similar to the original set of analyses.

Table S1. Mean numbers of stimuli per condition. Range across participants is given in brackets.

|  | Pattern estimator - Training set | Visible stimuli on tracking task | Imagined stimuli on tracking task |
| --- | --- | --- | --- |
| All trials | 959.81  [957-960] | 1432.40  [1394-1452] | 1467.20  [1425-1452] |
| Correct trials only | As above  (passive task) | 1240.20 [904-1408] | 1271.80 [956-1446] |

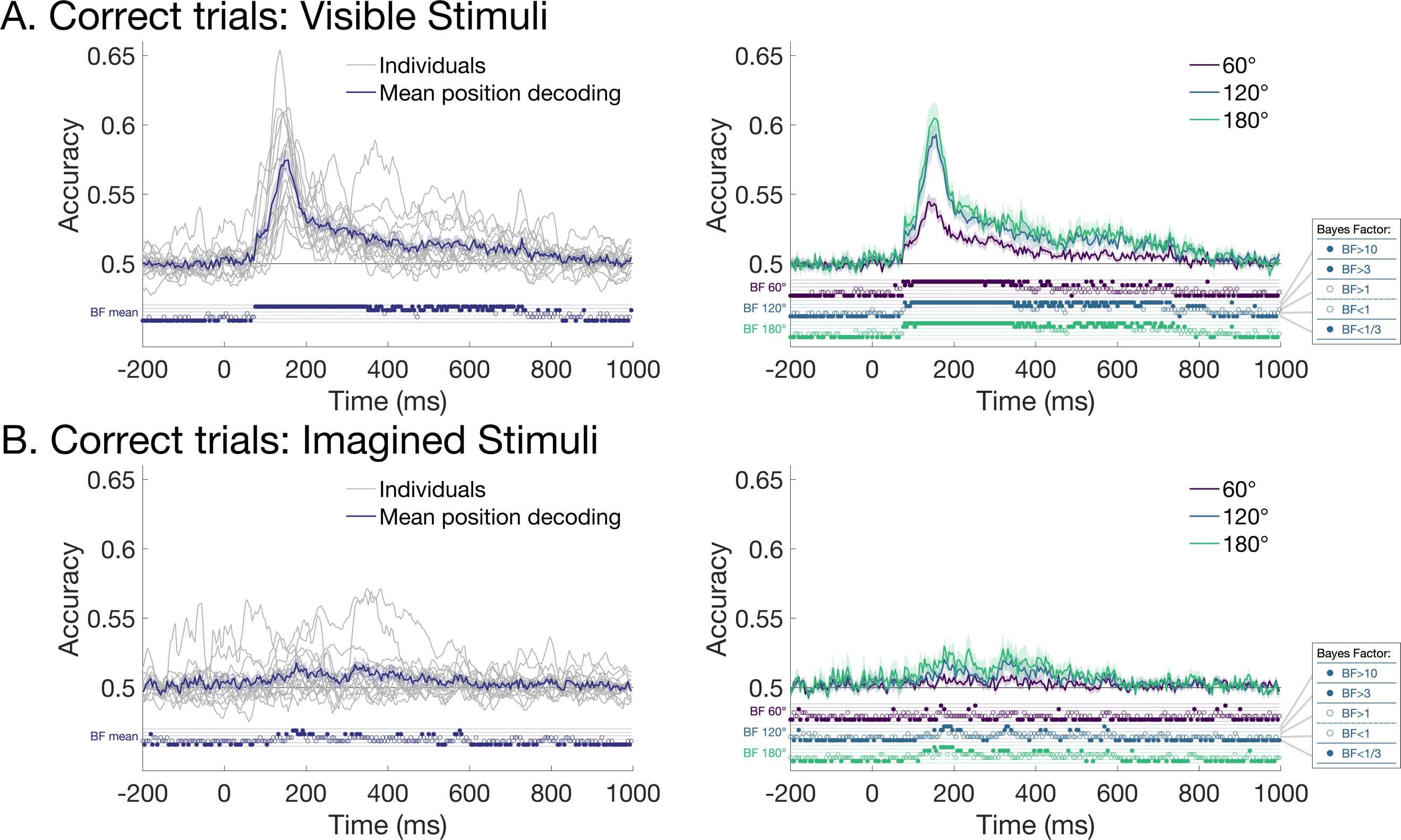

Figure S1. Decoding position using correct trials only. A. Visible stimuli. B. Imagined stimuli. The results are largely the same as the original decoding analyses which used all trials regardless of performance on the task.

**S2. Bayes Factors associated with position decoding**

Figure S2 shows the Bayes Factors associated with whole brain position decoding from Figure 4. For both visible and imagined stimuli, there was considerable evidence for position-specific representations in the neural signal.

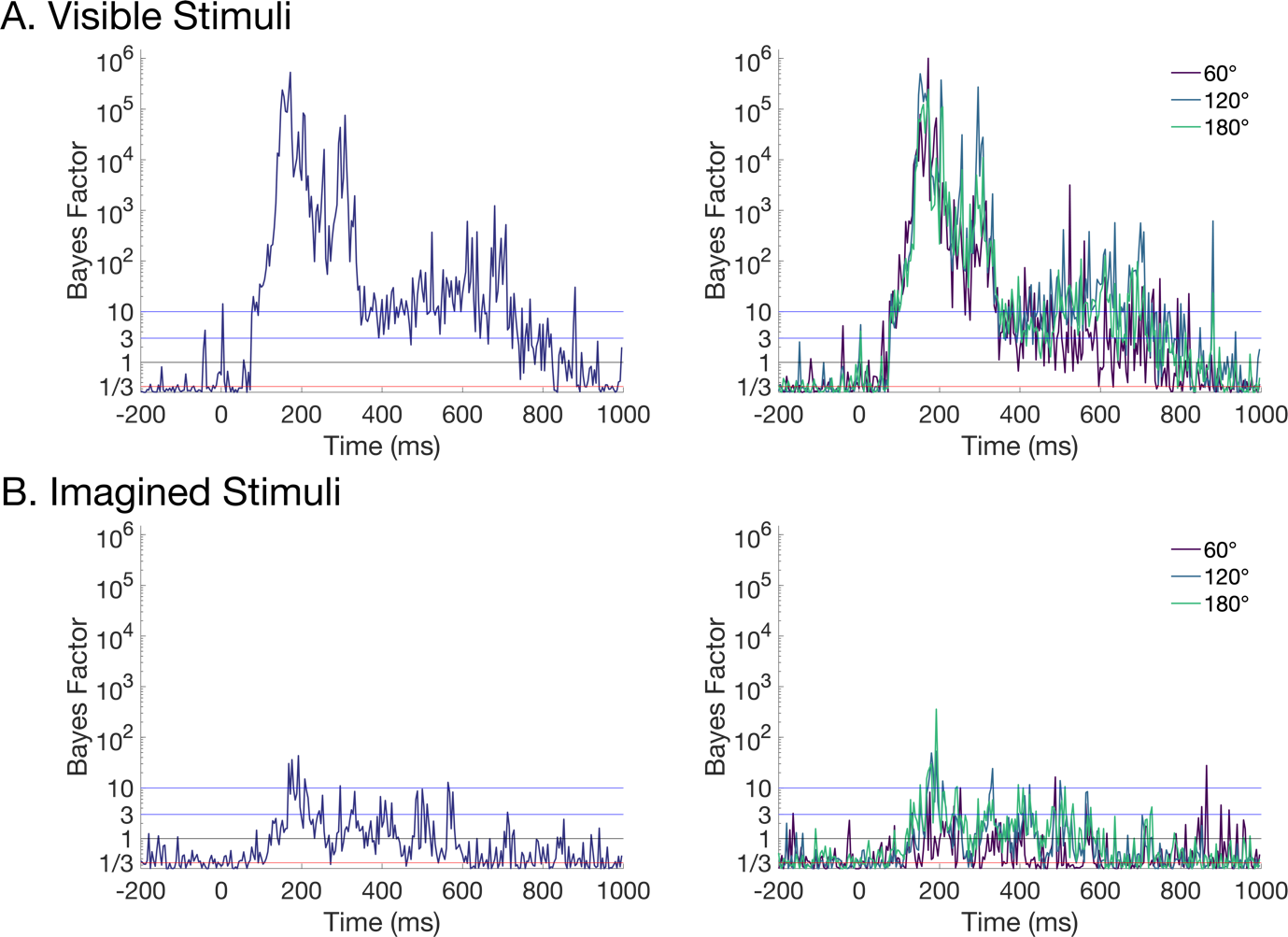

Figure S2. Bayes Factors (BFs) associated with position decoding accuracy as a function of time, plotted on log scale. A. Visible stimuli. B. Imagined stimuli. Left plots show BFs associated with decoding all stimulus positions, and right plots show BFs associated with decoding according to the angular distance between position pairs.

**S3. Analysis of position-related activity within frontal electrodes**

To assess the contribution of potential eye movements to the decoding results (and complement the posterior analysis), we performed decoding using a subset of electrodes from the front of the head. The 27 electrodes were Fp1, Fp2, AFz, AF3, AF4, AF7, AF8, Fz, F1, F2, F3, F4, F5, F6, F7, F8, FT7, FT8, FT9, FT10, FCz, FC1, FC2, FC3, FC4, FC5, FC6. Figure S3.1 shows decoding accuracy for classifiers trained and tested on the pattern estimator. Decoding was reliably above chance from approximately 150ms but considerably lower than whole brain analyses (Figure 3). For the cross-decoding analysis, when trained on the pattern estimator and tested on the tracking task, Figure S3.2 shows again that decoding is lower in general than the whole brain analysis. Furthermore, for imagined stimuli there is little evidence that the frontal electrodes contained position-specific representations (i.e., decoding is not reliably above chance except for a brief period just after 200ms).

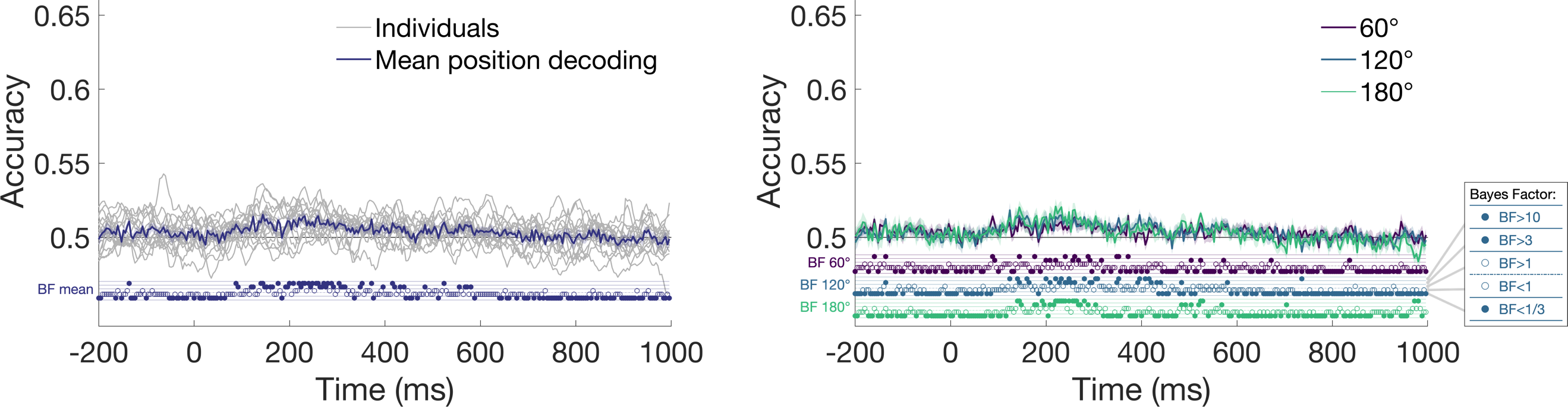

Figure S3.1. Position decoding over time for stimuli in the pattern estimator sequences using only frontal electrodes. Left plot shows mean position decoding and right plot shows position decoding as a function of the angular distance between stimuli.

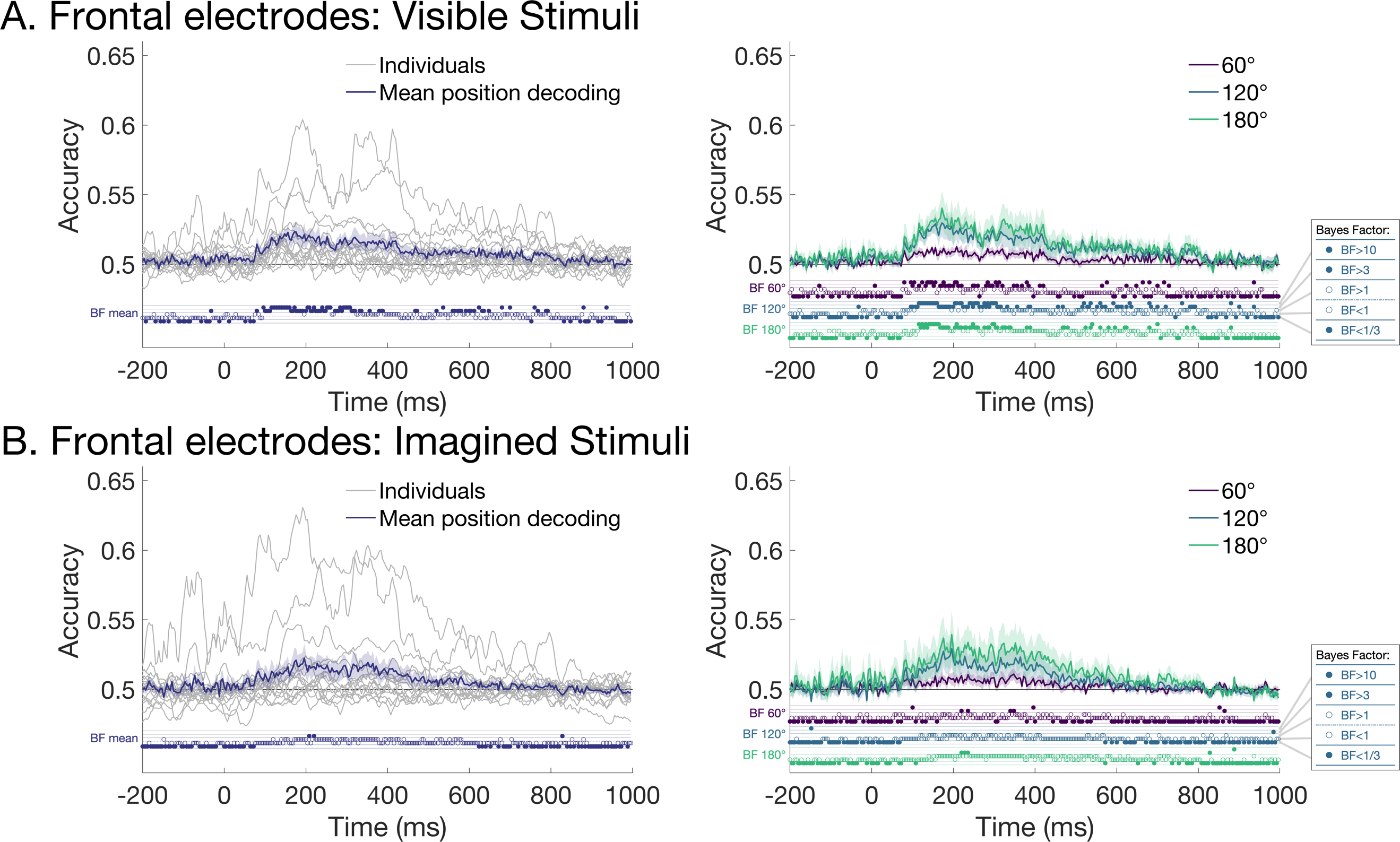

Figure S3.2. Decoding visible and imagined stimuli on the tracking task using only frontal electrodes. A. Visible stimuli. B. Imagined stimuli.

**S4. Decoding of stimuli in the tracking task by training and testing on the same condition**

All original analyses were performed by training classifiers on the pattern estimator sequences and testing on the tracking task. However, the cross-decoding analysis limits the results to information that is common to both types of experimental sequences. To assess position-specific information in the tracking task alone, we performed cross-validated leave-one-block-out decoding separately for the visible and imagined stimuli. Figure S4 shows that decoding accuracy was above chance for the whole time period, including prior to the stimulus being presented or imagined. Due to the predictable movement of the stimuli, above chance decoding prior to the stimulus could reflect anticipatory neural signals relating to the upcoming position, and/or decoding of the previous stimulus position.

The dynamics of the visible decoding looked qualitatively similar to, but higher than, the original cross-decoding analysis, with a peak around 150ms. Decoding of the imagined stimuli followed a different trajectory, with highest decoding at approximately 0ms, which was the time the tone was presented and when participants were meant to be imagining the stimulus position. Interestingly, imagined decoding resembled decoding in the cross-decoding time generalisation analysis (see Figure 6B), although again with higher decoding accuracy across the whole time period. It is impossible to make any strong claims about visible and imagined stimulus information based on these analyses because of the confounding positions of the previous and following stimuli. Nevertheless, it seems that the stimulus position information in the tracking task is not drastically different to stimulus position information in the pattern estimator, lending support to the idea that spatial imagery relies on stimulus-driven processes.

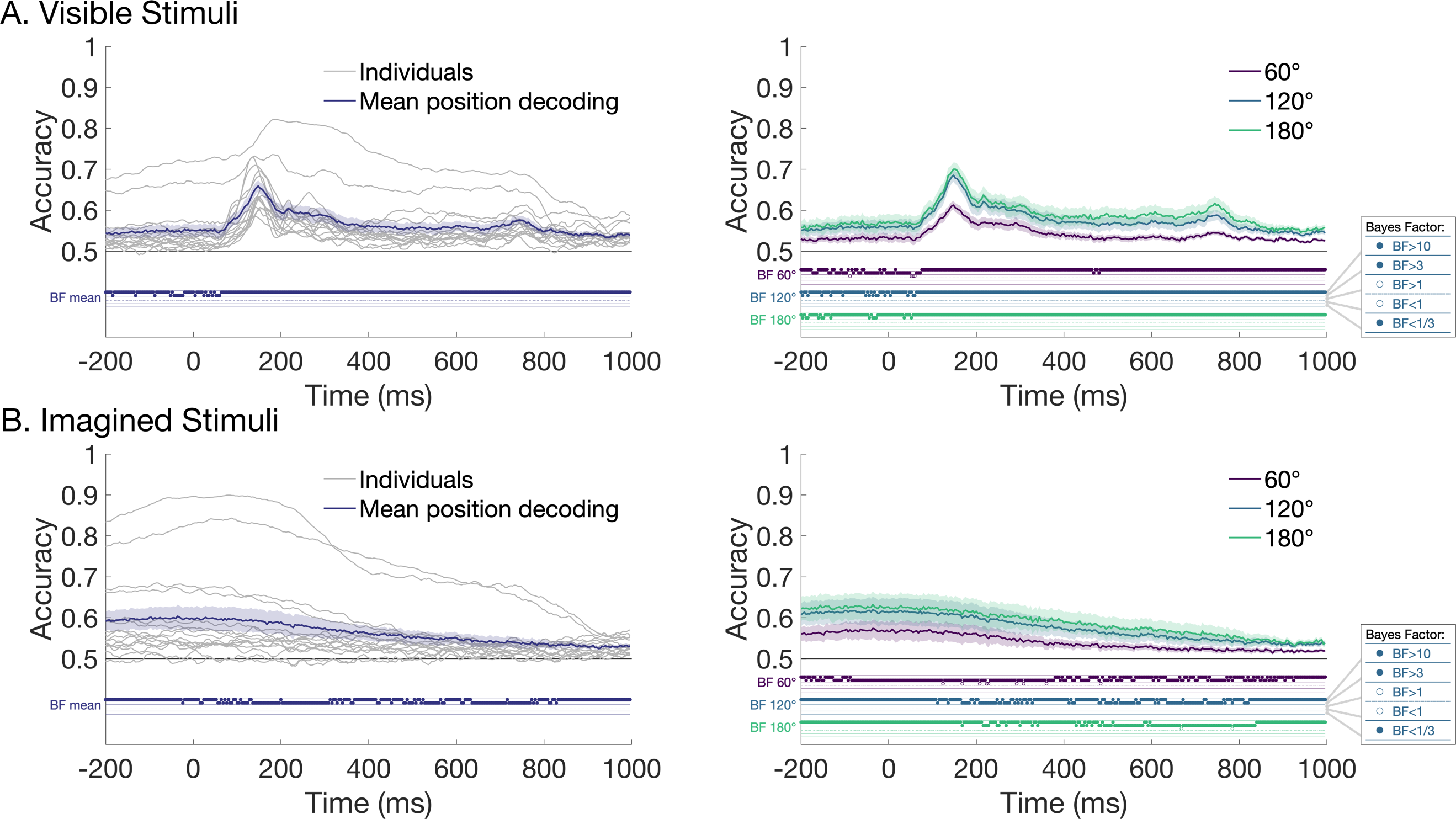

Figure S4. Decoding using leave-one-block-out cross-validation for stimuli on the tracking task. A. Visible stimuli. B. Imagined stimuli.

**S5. Neural responses to stimuli**

To assess the data quality and the temporal dynamics of neural activation for the different conditions in the study, we computed event-related potentials by averaging the signal from all relevant trials and electrodes over time, regardless of stimulus position (Figure S5.1). Data were preprocessed as in the decoding analysis (0.1Hz highpass, 100Hz lowpass, average reference). No trials were rejected. ERPs were calculated separately for frontal electrodes and posterior electrodes. For the pattern estimator, there were 960 trials per participant, except for one participant who only had 957 trials due to a data collection error. The number of trials was similar for visible stimuli (*M* = 1432.40, range 1394-1452) and imagined stimuli (*M* = 1467.20, range 1425-1475) per participant (see Table S1).

As can be seen in Figure S5.1, there was a clear event-related response for all stimuli. ERPs for the pattern estimator resembled a sine wave due to the rapid stimulus presentation. Dynamics for the slower tracking task look like characteristic event-related responses; for example, there was a positive peak at 100ms (P1 component) and negative peak at 180ms (N170 component) after every stimulus presentation. Imagined responses were slightly reduced relative to the visible stimuli, though note that no statistics were performed as this was a qualitative analysis. ERPs for each participant are shown in Figure S5.2.

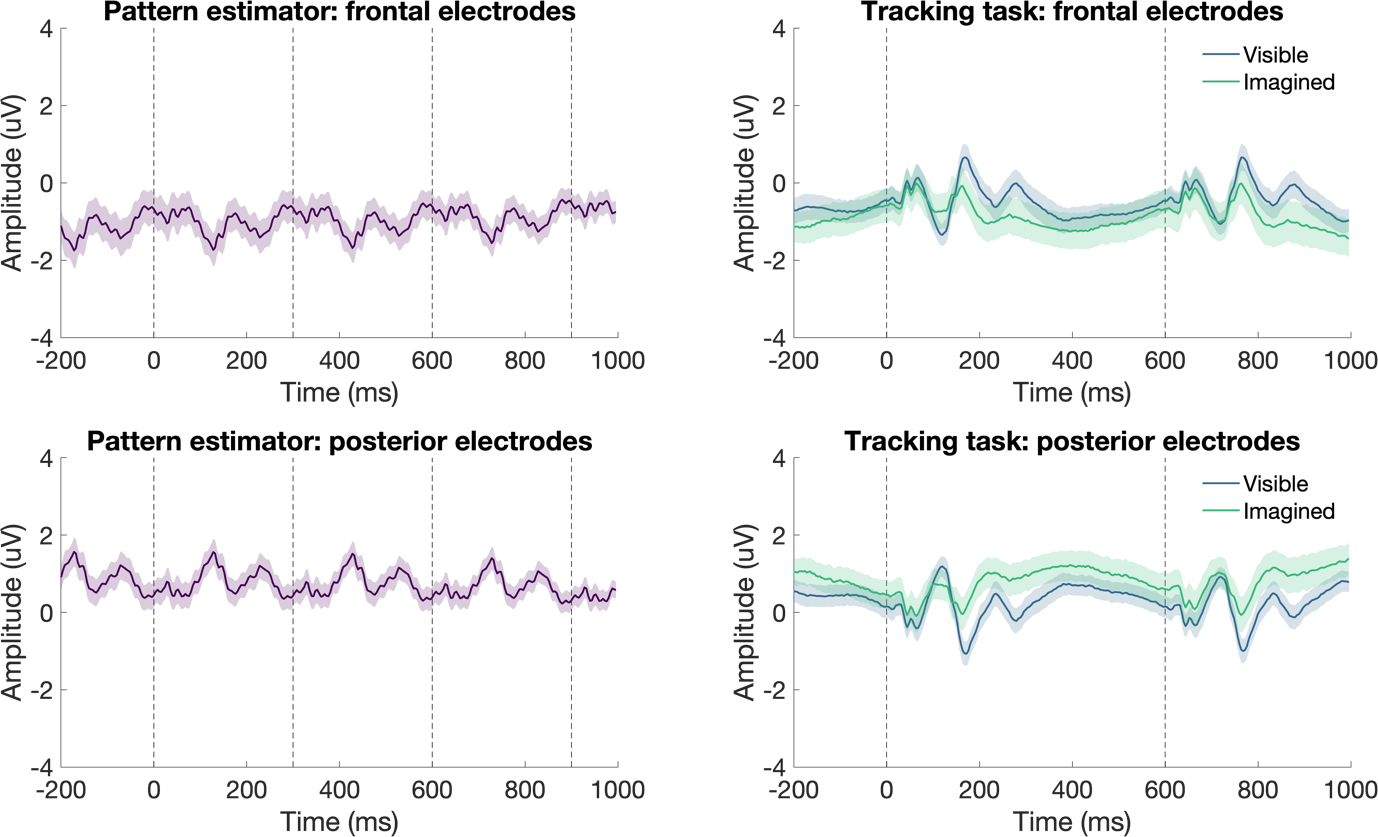

Figure S5.1. Event-related potentials for each condition and electrode cluster. Left plots show ERPs for stimuli in the pattern estimator sequences. Right plots show ERPs for visible and imagined stimuli in the tracking task. Dotted vertical lines denote onset of stimuli within the task. Shaded areas show standard error of the mean.

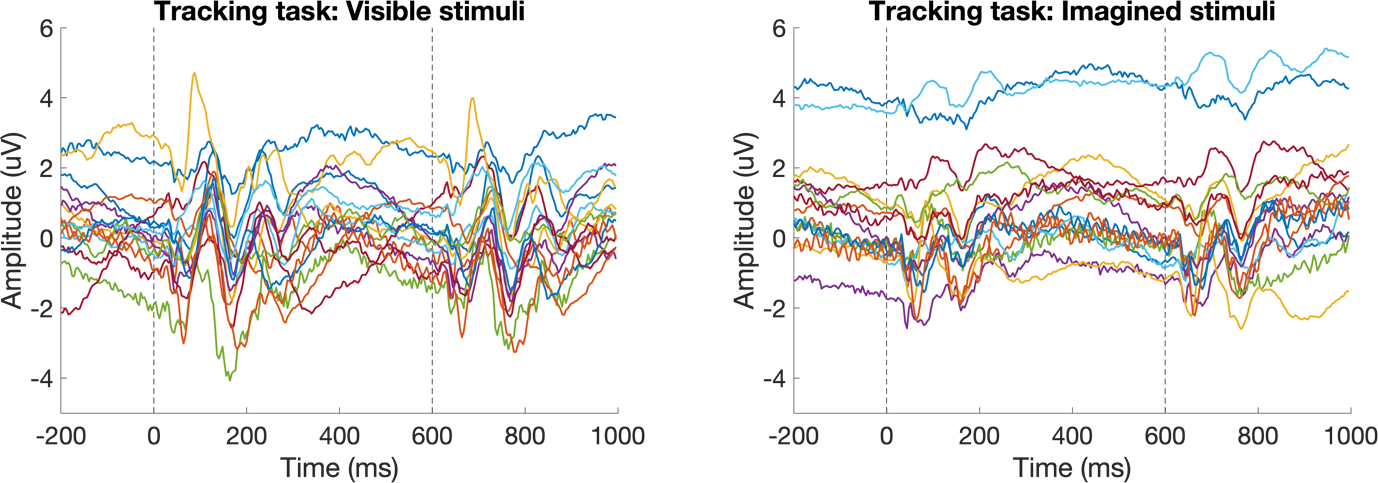

Figure S5.2. Event-related potentials from posterior electrode cluster. Left plots show visible ERPs and right plot shows imagined ERPs.

**S6. Individual participant decoding**

There was considerable variation in the magnitude of decoding accuracy across participants (see Figure S6). Imagined decoding showed more temporal variation across participants than visible decoding.

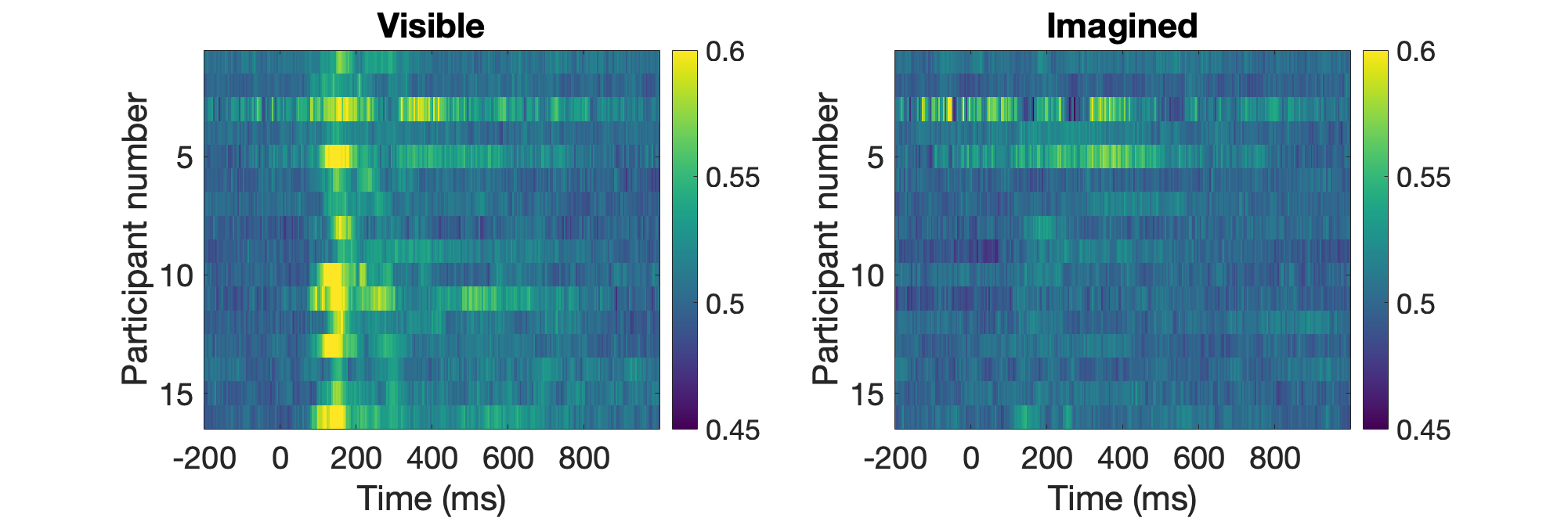

Figure S6. Decoding accuracy over time per participant for visible (left) and imagined (right) stimuli on the tracking task. Decoding models were trained on the pattern estimator.
